# Aging-induced hepatocyte CD44 drives IL6/STAT3 signaling and associates with impaired neighboring T cell function

**DOI:** 10.64898/2025.12.20.695732

**Authors:** Armin Gandhi, Kathryn Lande, Yichen Li, Filipe A. Hoffman, Michael LaPorte, Shirong Tan, Garrett Evensen, Benji Portillo, Marcos Garcia Teneche, Rouven Arnold, Adarsh Rajesh, Jessica Proulx, Shanshan Yin, Aaron Havas, Charlene Miciano, Qian Yang, Elizabeth Smoot, Sainath Mamde, Andrew Davis, Kevin Yip, Allen Wang, Bing Ren, April Williams, Susan Kaech, Peter D. Adams

**Author notes:** Corresponding authors: Peter D. Adams, Armin Gandhi.

## Abstract

Liver cancer incidences increase dramatically beyond 55 years of age, suggesting that age-associated changes contribute critically to tumor initiation. However, the mechanisms linking liver aging and cancer initiation are not well defined. This study investigates the role of CD44, a marker of liver tumor-initiating cells (TIC), in age-associated liver pathophysiology. Aged livers showed accumulation of CD44-expressing hepatocytes exhibiting enrichment of immune modulatory genes and activation of the immunosuppressive IL6/JAK/STAT3 pathway. Indeed, in adoptive transfer assays, antigen-exposed CD8+ T cells mounted a lower IFN-γ response in aged livers than in young livers, indicating an immunosuppressive aged milieu. Concordantly, spatial analyses showed that the proximal neighbourhoods of *Cd44*-expressing hepatocytes are enriched in T cells exhibiting reduced cytokine and chemokine gene expression. Finally, hepatocyte-specific knock out of *Cd44* mitigated the IL6/JAK/STAT3 gene signature in aged livers. Overall, these findings suggest that CD44 expression in aged hepatocytes promotes activation of the immunosuppressive IL6/JAK/STAT3 pathway and this is associated with impaired T cell effector function.

## 1. Introduction

Age is a major risk factor for hepatocellular carcinoma (HCC)^1,2^. In the US, most primary HCC incidences occur over 55 years of age^3^. Therefore, it is important to understand the role of aging in HCC initiation. Aging is associated with immune dysfunction characterized by low grade chronic inflammation and impaired immune surveillance^4^. Specifically, aging is characterized by decreased thymic output^5^ impaired T cell activation and proliferation^6^ and increased dysfunctional CD8+ T cells^7,8^. Such changes have been proposed to contribute to cancer initiation in aged individuals^9,10^. In the liver, early hepatic neoplasia have been shown to possess tertiary lymphoid structures with features of T cell activation, antigen presentation, as well as immunosuppression and exhaustion^11^, suggesting that although local immune activation occurs early in liver carcinogenesis, it may not be sufficient to prevent tumor initiation. Notably, Interleukin 6 (IL-6) known to mediate chronic inflammation in the liver^12,13^ and immune suppression in HCC^14,15^ increases with age^13^. IL-6 exerts it effects, at least in part, through JAK/STAT3 signaling. However, upstream determinants of IL6/JAK/STAT3 activity in aged liver are not well defined.

CD44 is a transmembrane glycoprotein receptor, primarily for hyaluronic acid, but has also been shown to interact with osteopontin, collagen and fibronectin. Moreover, CD44 is a coreceptor for EGFR and c-MET^16^ and participates in NF-κB and STAT3 activation^19,20^. CD44 is a well-established marker for cancer stem cells or tumor-initiating cells (TICs) in the liver^19–21^. Indeed, CD44 is not expressed on normal hepatocytes but is expressed on hepatocellular carcinoma progenitor cells (HcPCs) that arise early in DEN-treated livers^22^. *Cd44* expression is repressed by p53 in untransformed cells^23^. However upon liver damage, STAT3 causes induction of CD44 which then antagonizes p53 signaling leading to a pro-survival and pro-proliferative state enabling early tumor-initiating conditions^24^. In sum, CD44 has many reported functions that can potentially contribute to its role in liver cancer initiation. However, the role of hepatocyte CD44 in aged liver, where risk of HCC is increased, has not yet been defined.

Here, we investigated a potential role for hepatocyte CD44 in age-dependent changes to the liver that might contribute to HCC. Using single nucleus 10x multiome, spatial transcriptomics and hepatocyte-specific CD44 knockout, we find that CD44-expressing hepatocytes increase in aged male livers. This promotes upregulation of IL6/JAK/STAT3 pathway and is associated with features of local and tissue-wide suppressed T cell effector function.

## 2. Results

### 2.1. CD44-expressing hepatocytes increase in aged male livers

To begin to address the role of CD44 in aging, we first asked whether aging alters hepatocyte CD44 expression. RNA expression analysis from our previously published bulk RNAseq of isolated hepatocytes from young and old male mice^25^ revealed a significant increase in *Cd44* mRNA in 24-month-old liver hepatocytes as compared to hepatocytes from 5-month-old mice (Fig. 1A; median *Cd44* normalized expression in male hepatocytes: young = 58.17325, old = 131.2585; adjusted p value = 0.0145). Single nucleus RNA sequencing analysis of young and old male livers revealed a significant increase in *Cd44* transcript in hepatocytes in aged male livers (Fig 1B and S1A; mean *Cd44* expression Young = 0. 06292444, Old = 0.1501005; p value = 1.3×10^−14^). In addition, single nucleus ATAC sequencing further showed increased chromatin accessibility (which typically correlates with expression^26^) at the *Cd44* locus in aged male hepatocyte nuclei as compared to young (Fig. 1C; mean *Cd44* ATAC signal Young = 0. 189815, Old = 0.227709; p value = 1.2×10^−13^). CosMx spatial transcriptomics likewise showed that *Cd44* mRNA is higher in aged male livers as compared to young male livers (Fig. 1D and S1B). Finally, to confirm and quantify CD44 protein on the hepatocyte population, hepatocytes were isolated from young and old male livers and CD44 expression was quantitated by flow cytometry. Flow cytometry analysis revealed a significant increase in the percentage of CD44-expressing hepatocytes (gated on CD45-negative cells) in old male livers as compared to the young male livers (Fig. 1E, Fig. S1C, FigS1D; median young = 1.09%, median old = 2.95%). Single nucleus RNA sequencing analysis of young and old female livers revealed no difference in *Cd44* transcript in aged female hepatocytes (Fig S1E; mean RNA expression: Young: 0.05623523, Old: 0. 05118435). Altogether, by bulk and single cell RNA measures, and at the protein and chromatin levels, we observed a consistent increase in hepatocyte CD44 expression in aged male livers.

**Figure 1.**
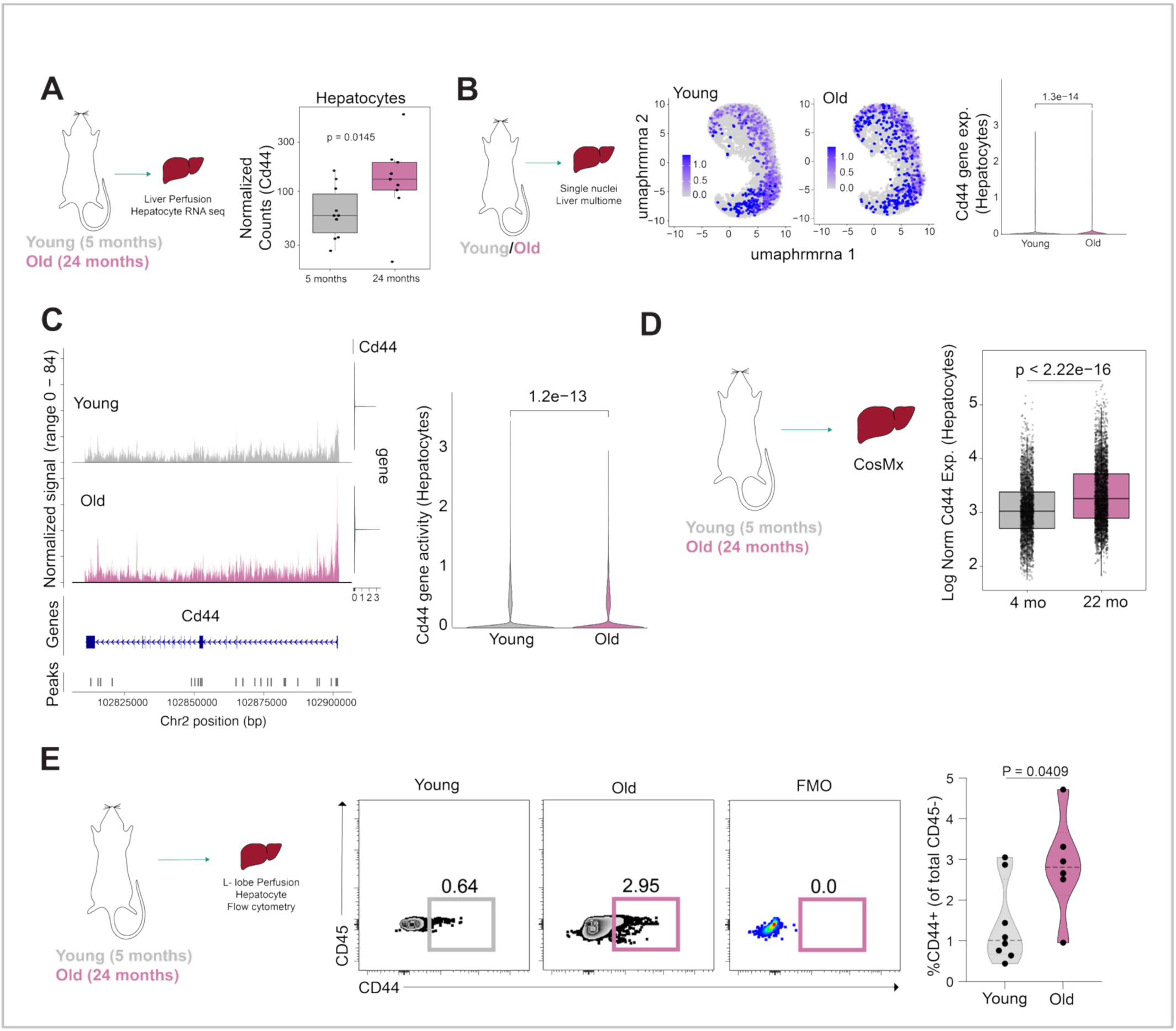
Abundance of CD44-expressing hepatocytes increases in aged mouse liver. **A.** Bulk RNA sequencing of young (5-months) and old (24-months) isolated hepatocytes^25^. Plot shows *Cd44* transcript levels as normalized counts (on Y-axis) per mouse. Each dot represents an individual mouse (n = 10 young, n = 9 old male livers). Adjusted p value was determined from DEG analysis with DESeq2, p = 0.0145. **B.** Single nucleus multiome profiling of young and old mouse livers. Uniform Manifold Approximation and Projection (UMAP) of nuclei from young (n = 3) and old (n = 3) male C57BL/6J mice annotated as hepatocyte cell type and showing *Cd44* transcript expression. Each dot represents a nucleus (young: 16441 nuclei, old: 10838 nuclei). Corresponding violin plots quantify *Cd44* expression across all hepatocyte nuclei from young and old livers. P value comparing young and aged was determined by Wilcoxon rank sum test, p = 1.3e-14. **C.** Coverage plot for ATAC seq signal at the *Cd44* locus from young and old hepatocytes (same livers as 1B). Corresponding violin plots quantify single nucleus ATAC seq signal at *Cd44* locus in all hepatocyte nuclei from young (n = 3) and old (n = 3) livers. P value comparing young and aged was determined by Wilcoxon rank sum test, p = 1.2e-13. **D.** CosMx spatial transcriptomic analysis of young (4-months) and old (22-months) liver sections. Sections were treated with H2O2. Box plot shows distribution of *Cd44* expression per hepatocyte with age, from 4 young and 5 old mice. Y-axis represents log-normalized expression of *Cd44* in hepatocytes. P values were determined by t test between log-normalized *Cd44* expression in young and old hepatocytes for all cells expressing *Cd44* (*Cd44*− cells were excluded), p < 2.22e-16. **E.** Flow cytometry analysis of CD44 in hepatocytes isolated from young and old mouse livers. L-lobes were perfused, and hepatocytes were purified using Miltenyi perfusion kit. Representative flow plots show CD44 versus CD45 staining with FMO (fluorescence minus-one) control indicated. Corresponding violin plot quantifies the percentage of CD44+ population within the CD45 negative population by age; each dot represents one mouse (n = 8 young, n = 6 old); p = 0.0409 was calculated by Unpaired t test with Welch’s correction.

### 2.2. *Cd44* expressing hepatocytes in aged male livers exhibit an immune modulatory gene signature enriched in IL6/JAK/STAT3 signaling

Given this increased abundance of CD44-expressing hepatocytes in livers of aged male mice and their known role in tumor initiation^22–24^, we set out to better define these aged CD44-expressing hepatocytes. Single nucleus RNA-seq (10x multiome) was used to identify differentially expressed genes (DEGs) comparing *Cd44* expressing (*Cd44*+) hepatocytes and *Cd44* non-expressing (*Cd44*−) hepatocytes in both young and old male mice livers. This differential expression analysis identified 4908 DEGs between *Cd44*+ and *Cd44*− hepatocytes in young mice (Fig. S2A; Upregulated in *Cd44+* = 1592, Downregulated in *Cd44+* = 3322 with Log2FoldChange cutoff of 0.1 and −0.1 respectively, and adjusted p-value ≤ 0.05, Supplementary Data T1) and 462 DEGs between *Cd44*+ and *Cd44*− hepatocytes in old mice (Fig. 2A; Upregulated in *Cd44+* = 305, Downregulated in *Cd44+* = 157 with Log2FoldChange cutoff of 0.1 and −0.1 respectively, and adjusted p-value ≤ 0.05, Supplementary Data T2). Interestingly, 255 DEGs (186 upregulated and 69 downregulated in *Cd44+*) were unique to the comparison of *Cd44*+ and *Cd44*− hepatocytes in old (Fig. 2B). Gene Ontology (GO) analysis of the 186 upregulated genes unique to old *Cd44*+ hepatocytes revealed a subset of terms related to antigen presentation and T cell interactions, specifically TAP1/2 binding, T cell receptor binding, peptide antigen binding (Fig. 2C, Supplementary Data T3). In addition, clustering of DEGs across all 4 hepatocyte populations (young *Cd44*−, young *Cd44*+, old *Cd44*− and old *Cd44*+) confirmed a cluster of upregulated DEGs (Cluster 2) enriched in old *Cd44*+ hepatocytes (Fig. 2D). GO term analysis of Cluster 2 also suggested immune modulatory gene signatures related to innate immune responses, cytokine signaling and antigen processing and presentation (Fig. S2B and Supplementary Data T4). More specifically, an IPA upstream regulator analysis of DEGs comparing old *Cd44*+ vs old *Cd44*− hepatocytes predicted activation of inflammation-related regulators such as IL6 and STAT3 in old *Cd44+* hepatocytes (Fig. 2E, Supplementary Data T5), key components in the IL6-JAK-STAT3 pathway that are frequently activated in cancer, including liver cancer^27,28^. The ATAC-seq data of our single nucleus multiome data confirmed increased accessibility of STAT3 transcription factor binding sites in old *Cd44*+ hepatocytes (Fig. 2F). We next used CosMx and MERSCOPE spatial transcriptomics to validate activation of the IL6-JAK-STAT3 pathway detected in our single nucleus multiome data. Using CosMx and MERSCOPE, *Cd44*+ hepatocytes were defined as those with normalized mRNA expression of *Cd44* > 0. Consistent with the single nucleus multiome data, we observed the highest expression of IL6-JAK-STAT3 upregulated genes^29^ in aged *Cd44*+ hepatocytes by both MERSCOPE (Fig. 2G) and CosMx (Fig. 2H), compared to old *Cd44*−, young *Cd44*+ and young *Cd44*−. In both cases, expression of *Cd44* itself served as internal validation. For MERSCOPE, a panel of control genes^30^ confirmed the specificity of the increase in IL6-JAK-STAT3 gene signature in old *Cd44*+ hepatocytes (Fig. S2C). As further validation, GSEA of mRNA sequencing from young and old isolated hepatocytes^25^ revealed significant positive enrichment of the IL6/JAK/STAT3 gene set in old hepatocytes (Enrichment score = 0.60, p = 2.14 × 10⁻³) (Fig. S2D). Together, these results show that *Cd44* expressing hepatocytes acquire an altered transcriptome modified with age to include immune modulatory signatures, notably activation of the IL6/JAK/STAT3 pathway.

**Figure 2.**
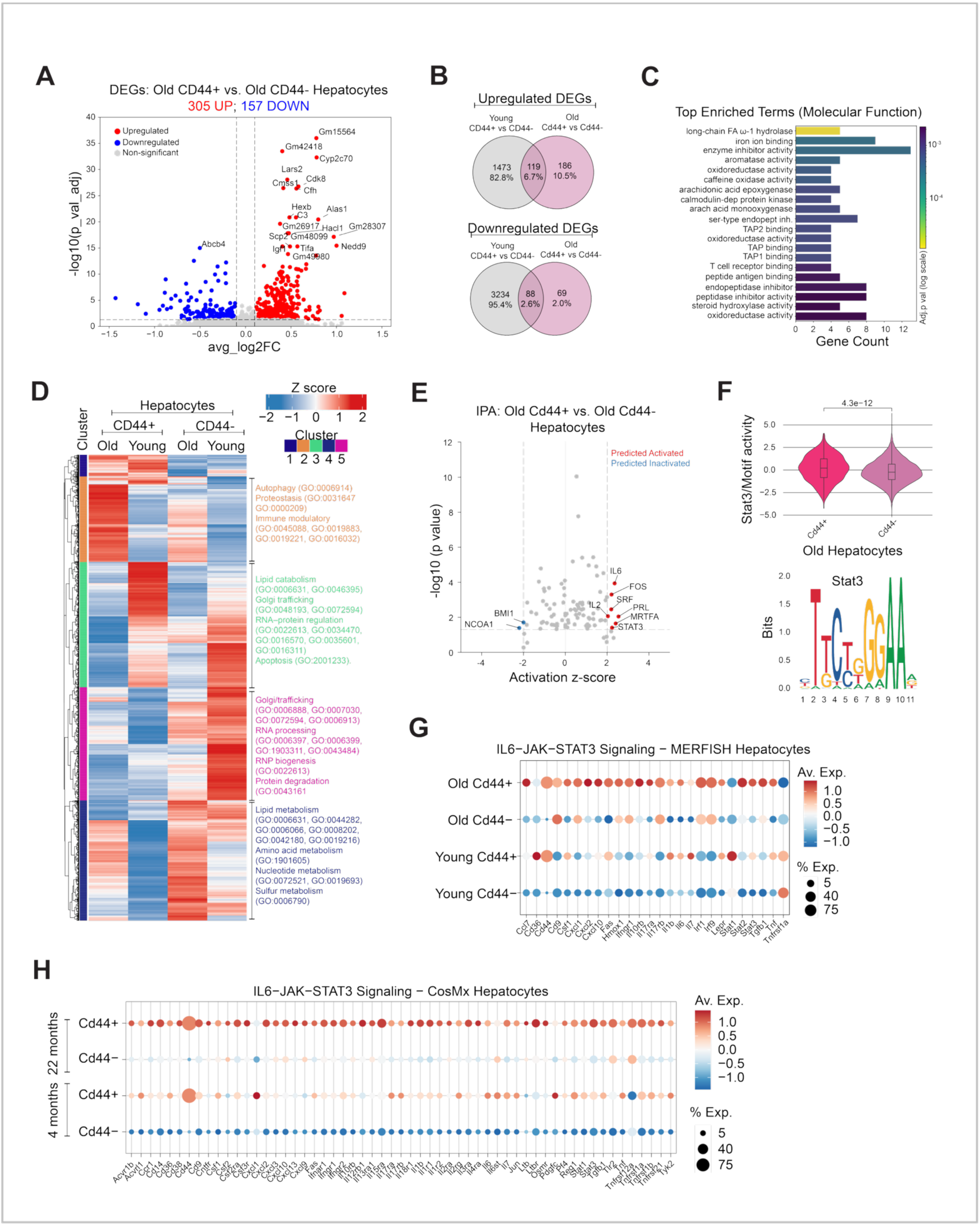
*Cd44*-expressing hepatocytes in aged livers exhibit an immunomodulatory transcriptome. **A.** Volcano plot depicting differentially expressed genes between *Cd44*+ versus *Cd44*− hepatocytes in old male livers. 305 genes are upregulated and 157 genes are downregulated in old *Cd44*+ hepatocytes (Log2FoldChange cutoff of 0.1 and −0.1 respectively, and adjusted p-value ≤ 0.05). Red = upregulated, blue = downregulated, grey = non-significant. **B.** Venn diagrams showing overlaps of upregulated and downregulated DEGs from *Cd44*+ versus *Cd44*− hepatocytes from young and old livers. 186 upregulated and 69 downregulated genes are unique to old *Cd44*+ hepatocytes. **C.** Top enriched GO terms obtained by analyzing 186 genes that are uniquely upregulated in old *Cd44*+ hepatocytes. **D.** DEGs by single nucleus RNAseq comparing *Cd44*+ versus *Cd44*− hepatocytes from all livers (n = 3 young and n = 3 old). Heat map shows unsupervised clustering of mean DEG expression (n=3 livers) in young and old, *Cd44*+ and *Cd44*− hepatocytes with summarized GO terms for major clusters. **E.** Ingenuity pathway analysis of DEGs comparing *Cd44*+ versus *Cd44*− hepatocytes in old liver. Volcano plot depicts top upstream predicted activated (Activation z-score ≥ 2) and predicted inactivated (Activation z-score ≤ −2) pathways in old *Cd44*+ hepatocytes. **F.** STAT3 Motif activity in old *Cd44*+ and *Cd44*− hepatocytes. Motif activity for all cells was calculated using ChromVar. Motif activity in old *Cd44*+ hepatocytes was compared to old *Cd44*− hepatocytes. P value was determined using Wilcoxon rank sum test, p = 4.3e-12. **G.** Expression analysis by MERSCOPE. Plot shows expression of IL6/JAK/STAT3 pathway genes in old and young *Cd44*+ and *Cd44*− hepatocytes. Color denotes average expression (n = 2 young, n = 2 old) and size denotes % expressing cells. **H.** Expression analysis by CosMx. Plot shows expression of IL6/JAK/STAT3 pathway genes in old and young *Cd44*+ and *Cd44*− hepatocytes. Color denotes average expression (n = 4 young, n = 5 old) and size denotes % expressing cells.

### 2.3. *Cd44*-expressing hepatocyte neighborhoods are associated with cytokine- and chemokine-deficient T cells in aged livers

Expression of STAT3 in tumor cells is known to drive an immunosuppressive microenvironment through secreted mediators such as IL-6 and IL-10^31^. In addition, IL-6 is known to inhibit IFN-γ and TNF-α in activated T cells^32^. Considering these observations, we asked whether the aged hepatic environment dampens antigen-specific function of otherwise young, healthy CD8+ T cells. To do this, we expressed LCMV GP33-41 (KAVYNFATC) antigen in hepatocytes in young and old mice by retro-orbital injection of AAV8-TBG-GP33 (or AAV8-TBG-MCS as empty vector control). Two weeks later, *ex vivo* activated GP33-specific P14 CD8+ T cells (from a young donor P14 mouse^33,34^) were injected into young and old GP33 (or MCS control) expressing mice and mice were culled 5 days later. As expected, P14 T cells accumulated comparably in the livers of young and old mice that express GP33 but not in control livers or the spleen (Fig. S3A for gating strategy, S3B). GP33 antigen encounter was comparable between young and old mice based on PD-1 expression on P14 CD8+ T cells (Fig. S3C). However, IFN-γ expression was significantly lower in P14 CD8+ T cells recovered from old recipient livers versus young livers (Fig. 3A), suggesting dampening of antigen-specific T cell activation in aged livers.

**Figures 3.**
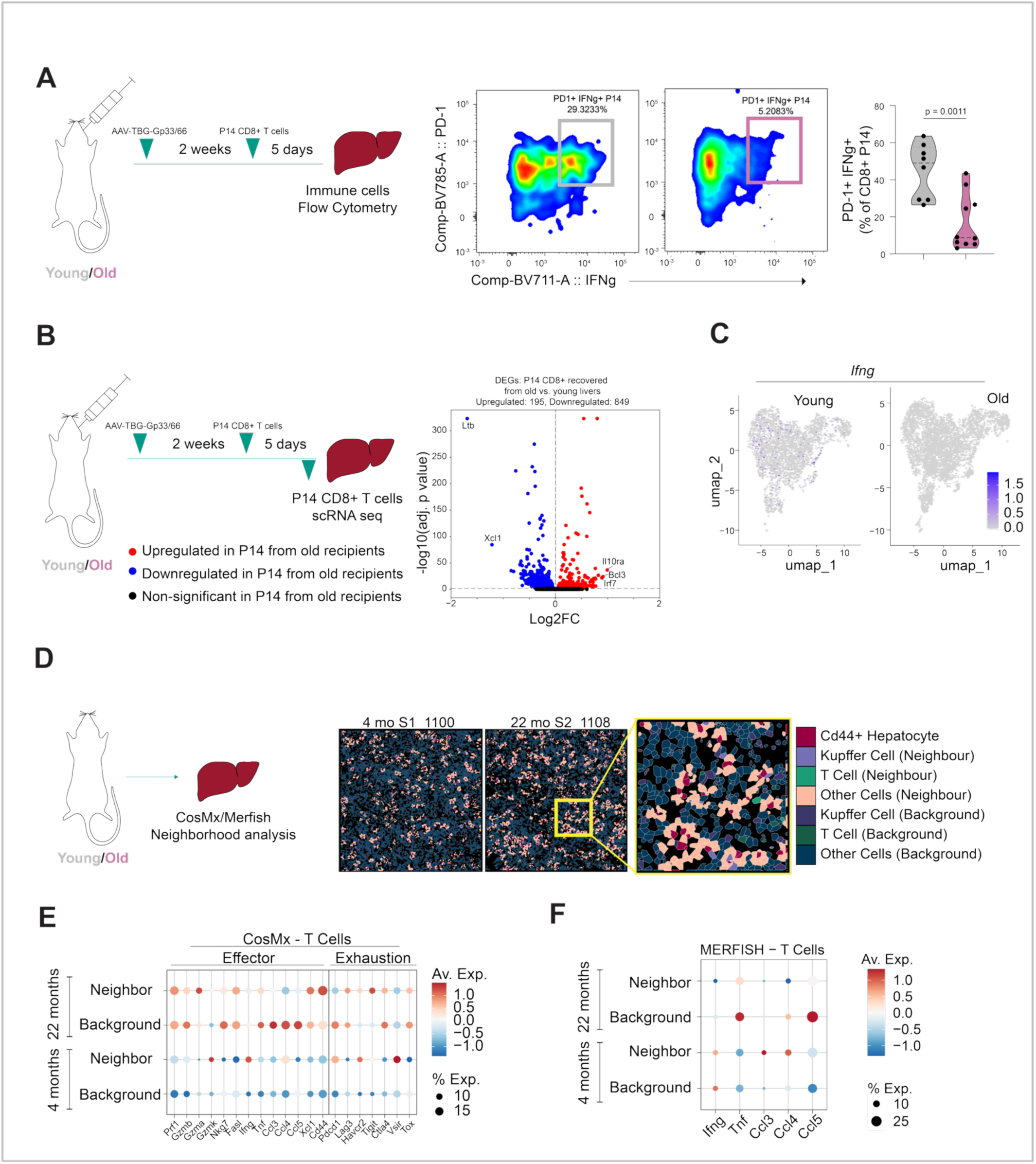
Aged livers and *Cd44*-expressing hepatocyte neighborhoods in aged livers harbor T cells with markers of inactivation. **A.** Flow cytometry analysis of recovered P14 CD8+ T cells from GP33-expressing young and old livers. Representative flow plots show IFN-γ versus PD-1 staining. Corresponding violin plot quantifies the percentage PD-1+, IFN-γ+ of the CD8+ P14 population recovered from young and aged livers; each dot represents one mouse (n = 8 young, n = 10 old); p value was determined by Unpaired t test with Welch’s correction, p = 0.0011. **B.** Single cell RNA seq analysis of P14 CD8+ T cells recovered from GP33-expressing young and old livers. Cells pooled from n = 7 young and n = 7 old mouse livers. Volcano plot shows DEGs comparing P14 CD8+ T cells recovered from old versus young livers. Significant ( p < 0.05) DEGs with a log2FC of more than 1 or less than −1 have been labelled, red = upregulated, blue = downregulated, black = non-significant. **C.** UMAP shows *Ifng* expression in P14 CD8+ T cells recovered from young and old livers, each dot represents a cell, all cells were pooled from 7 young and 7 old mouse livers. **D.** CosMx analysis of young and old livers. Representative spatial images showing *Cd44*+ hepatocytes and the cells analyzed as neighbors and background in young and old liver sections. **E.** Effector and exhaustion marker expression in neighbor and background T cells of young and old livers by CosMx. Color denotes average expression (n = 4 young, n = 5 old) and size denotes % expressing cells. **F.** Effector cytokine (*Ifng* and *Tnf*) and chemokine (*Ccl3*, *Ccl4*, *Ccl5*) expression in neighbor and background T cells of young and old livers by MERSCOPE. Color denotes average expression (n = 2 young, n = 2 old) and size denotes percent expressing cells.

To determine more broadly how age affects the transcriptome of P14 CD8+ T cells, we performed single cell RNA-seq analysis of the P14 CD8+ T cells recovered from young and old livers. Confirming the immunostaining (Fig. 3B), P14 T cells from old livers expressed less *Ifng* mRNA (Fig. 3C). Differential expression analysis revealed 195 upregulated genes and 849 downregulated genes (p < 0.05 and log2FC of greater than or less than 0) in the recovered P14 CD8+ T cells from old livers compared to young livers (Fig. 3B; Supplementary Data T6). GO enrichment analysis of the significant DEGs confirmed downregulation of genes related to *Ifng* production in P14 CD8+ T cells recovered from old livers compared to young, and also downregulation of genes related to T cell chemotaxis and migration (Fig. S3D). Consistent with decreased *Ifng* expression, we also observed a decrease in the effector cytokine *Tnf* and chemokines (*Ccl3* and *Ccl4*) in P14 T cells from old mice (Fig. S3E). Upstream regulator analysis of the DEGs also predicted reduced activity of MYC and IL2, important for CD8+ T cell function^35–37^ in the P14 CD8+ T cells recovered from old livers (Fig. S3F, full list in Supplementary Data T7). Likewise, GSEA revealed significant downregulation in IL2/STAT5 pathway related genes in P14 CD8+ T cells from aged livers (Fig. S3G). Overall, these findings confirmed that the aged milieu alters antigen-specific CD8+ T cells, most notably by reducing expression of *Ifng*, *Il2* and *Tnfa*.

Given that we previously observed that old *Cd44*+ hepatocytes are characterized by high IL6/JAK/STAT3 pathway related genes, we next asked whether *Cd44*-expressing hepatocytes are associated with impaired effector functions in neighboring T cells. To address this question, we mined CosMx and MERSCOPE data to assess the spatial relationship between *Cd44*+ hepatocytes and markers of T cell activation and exhaustion in old liver. *Cd44*+ hepatocyte neighbors were defined by ranking all cells by their physical proximity to each *Cd44*+ hepatocyte using a K-nearest neighbor algorithm (K = 8). (Fig. 3D). Regions outside of these defined neighborhoods were annotated as background. Cell type labels were assigned based on canonical marker expression (Fig. S3H and S3I). Aligning with the single cell RNA-seq of P14 T cells (Fig 3C and S3E), CosMx analysis revealed lower effector cytokine (*Ifng*, *Tnf*) and chemokine (*Ccl3*, *Ccl4*, *Ccl5*) expression in T cells of the aged *Cd44*+ neighborhoods, as compared to T cells in the aged background (Fig. 3E), and this was validated by MERSCOPE (Fig. 3F). This gene expression pattern in T cells was specific to old neighborhoods versus background and not observed for young neighborhoods versus background. Cytotoxicity associated transcripts showed no consistent difference, except for *Nkg7* which was lower in aged neighboring T cells, by both CosMx and MERSCOPE. Overall, spatial analysis across CosMx and MERSCOPE indicated that aged *Cd44*+ hepatocyte neighborhoods contain T cells exhibiting reduced effector cytokines and chemokines, suggesting that aged *Cd44*+ hepatocytes may impair the function of T cells in their neighborhood. Altogether, the aged liver microenvironment suppressed antigen-specific CD8+ T cell effector cytokine pathways and in aged liver *Cd44*+ hepatocytes are associated with a local milieu of suppressed T cells.

### 2.4. CD44 depletion attenuates immune modulatory gene signatures in aged male mouse livers

Together, these results suggest that *Cd44*-expressing hepatocytes in aged livers are enriched for inflammatory and immune suppressive IL6/JAK/STAT3 signaling and are spatially associated with functionally impaired T cells within the immune suppressive aged liver. To test the causal role of CD44, we assessed the consequences of *Cd44* inactivation in aged liver. To do this, we knocked out *Cd44* in aged hepatocytes *in vivo* by retro-orbital injection of AAV8-TBG-saCAS9-U6-sgRNA-Cd44, an approach previously described by Zhang and coworkers^38^. We employed two different sgRNAs targeting *Cd44* (guide 2 and guide 4) and an sgRNA targeting Rosa26 as control. 8 weeks later, knockout efficiency was validated by guide RNA-directed genomic DNA sequencing which showed 22% knockout score with guide 2 and 35% knockout score with guide 4 (Fig. S4A and S4B). We analyzed guide 4 infected livers to validate CD44 depletion at the protein level by flow cytometry (Fig. S4C, S4D and S4E). To assess whether CD44 depletion alters age-associated inflammatory pathways, we performed RNA sequencing of whole livers from hepatocyte-specific *Cd44* knockout and control animals. Principle component analysis (PCA) revealed that old *Cd44* KO samples clustered with old Rosa26 control livers (Fig. 4A), suggesting a modest overall effect of *Cd44* KO. However, unbiased GSEA of transcriptomes from hepatocyte *Cd44* knockout versus controls revealed significant enrichment of metabolic pathways alongside concurrent suppression of ECM/adhesion, epithelial secretory, and immune/inflammation-dominated gene sets (Fig. 4B). More specifically, GSEA showed that *Cd44* KO resulted in a significant or trend to a decrease in some inflammatory pathways, including the IL6/JAK/STAT3 pathway (Fig 4C, 4D).These transcriptome-wide shifts are consistent with relief of chronic inflammatory tone in aged male livers suggesting that CD44-expressing hepatocytes contribute to inflammatory IL6/JAK/STAT3 signaling in aged male livers.

**Figure 4.**
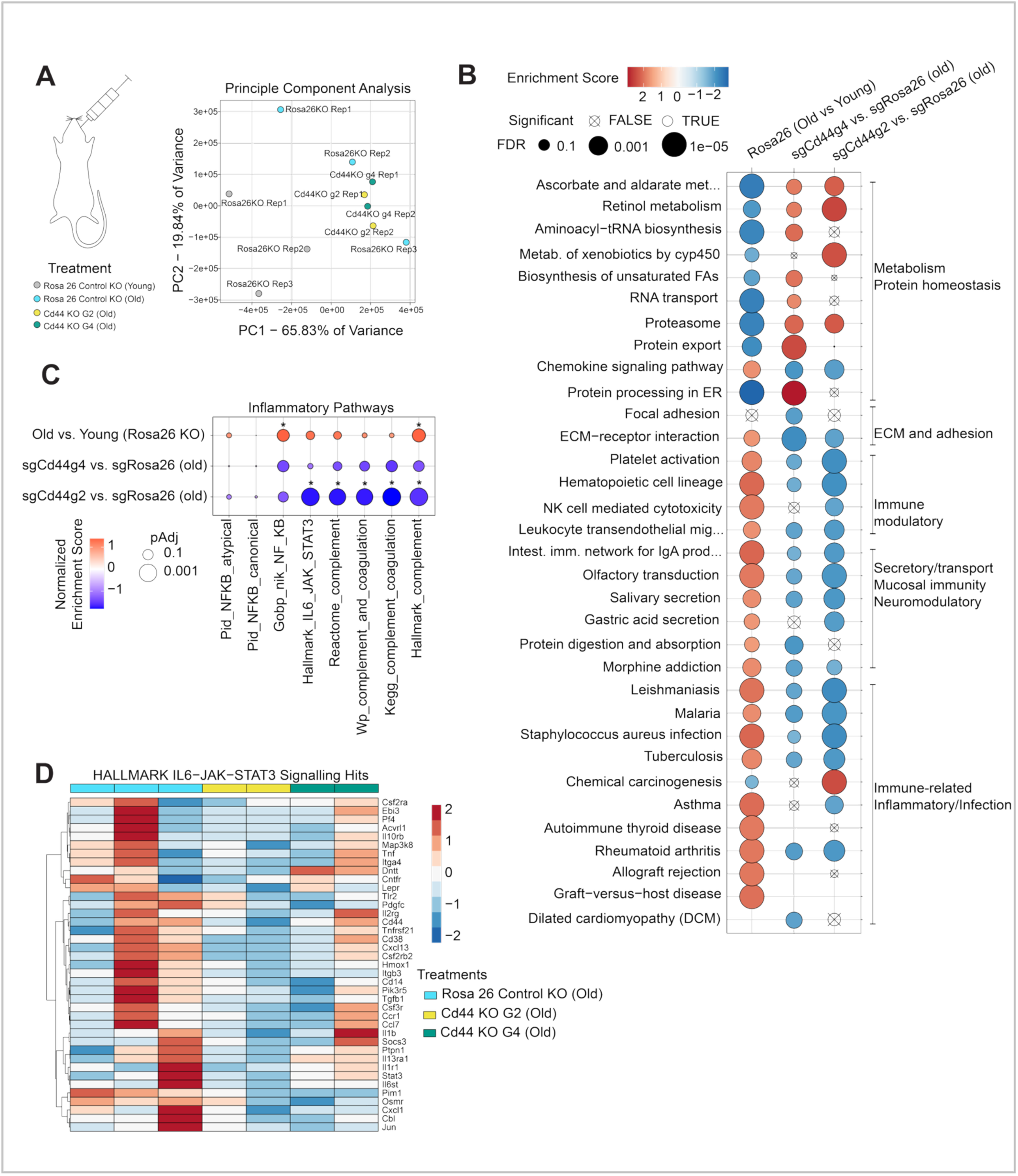
Knock out of hepatocyte *Cd44* attenuates immune modulatory gene signatures in aged mouse livers. **A.** RNAseq analysis of young and old control *Rosa26* sgRNA liver and *Cd44* knockout (with guides sgRNAg2 and sgRNAg4) liver. PCA plot shows transcriptome differences between livers. Each dot represents a mouse liver. N = 3 young and old Rosa26 control guide RNA, N = 2 Cd44 g2 and N = 2 Cd44 g4 old. **B.** Unbiased GSEA for DEGs comparing old *Rosa26* versus young *Rosa26* control and old *Cd44* knockout (g2 and g4) versus old *Rosa26* control. Color indicates normalized enrichment score (red = positive enrichment, blue = negative enrichment), and size of dot indicates adjusted p value. **C.** GSEA showing NFκB, IL6/JAK/STAT3 and complement/coagulation pathways of DEGs comparing old *Rosa26* versus young *Rosa26* sgRNA controls and old *Cd44* knockout (g2 and g4) versus old *Rosa26* controls. Color indicates normalized enrichment score (red = positive enrichment, blue = negative enrichment), and size of dot indicates adjusted p value. Significant p values are denoted by *; adjusted p < 0.05. **D.** Heatmap of scaled median of ratios normalized counts of genes in the leading edge of the g2 vs. old Rosa26KO or g4 vs. old Rosa26KO GSEAs for Hallmark IL6/JAK/STAT3 pathway. Rosa 26 control (n = 3), Cd44g2 (n = 2) and Cd44g4 (n = 2) old livers.

## 3. Discussion

Overall, our findings reveal a novel role of CD44 expression in hepatocytes in aging. We observed that (i) the abundance of CD44-expressing hepatocytes increases in aged male liver; (ii) Cd44-expressing hepatocytes show enrichment in immune modulatory gene ontologies and activation of the IL6/JAK/STAT3 pathway signature; (iii) the aged liver milieu reduced markers of antigen-specific CD8+ T cell function, specifically expression of *Ifng* and *Tnfa* transcripts and the IL2/STAT5 pathway signature; (iv) the neighbourhood of *Cd44*-expressing hepatocytes in aged liver is enriched in T cells with reduced *Ifng* and *Tnfa* gene expression; (v) hepatocyte-specific Cd44 knockout caused transcriptomic attenuation of the IL6/JAK/STAT3 pathway signature genes in the aged whole liver. Together, these observations implicate age-associated expression of hepatocyte CD44, known for its role in liver cancer initiation^24,26^, in activation of the immune suppressive IL6/JAK/STAT3 pathway and T cell dysfunction in the aged liver.

Prior work shows that CD44 induction antagonizes p53-mediated signaling to promote a pro-survival state^25, 26^. Our data show that *Cd44*-expressing hepatocytes are associated with T cell dysfunction and an immune suppressive milieu. Moreover, CD44 promotes expression of the IL6/JAK/STAT3 pathway signature, a pathway known to be immune suppressive through direct actions of IL6 on CD8+ T cells^32^ or through indirect mechanisms involving regulatory T cells and innate immune mediators^39^. Across two different spatial modalities, we observed that surrounding *Cd44* expressing hepatocytes in aged liver there was a consistent decline in T cell expression of *Ifng*, important for anti-tumor immunity^40^. Age-associated attenuation in immune surveillance may help in persistence of CD44 expressing hepatocytes, reinforcing the pro-oncogenic niche. Thus, our data support a model in which age-associated CD44 expression in hepatocytes contributes to a HCC-prone niche by fostering an IL6/JAK/STAT3-dependent immune suppressive microenvironment.

The mechanism underlying CD44 expression and the causal link between CD44 and IL6/JAK/STAT3 in aged hepatocytes is not known. However, CD44 is known to be induced by STAT3 activation^24^, and STAT3 is activated by CD44^18^. These observations point to a positive feedback loop between CD44 and STAT3 signaling, suggesting that sustained activation of both CD44 and STAT3 may be important for persistence of the pre-cancerous hepatocyte state. Accordingly, targeting age-associated CD44 in hepatocytes may be a potential strategy to modify early oncogenic risk and suppress HCC. As partial proof-of-principle, we showed that hepatocyte-specific Cd44 knockout dampened the IL6/JAK/STAT3 pathway signature genes. However, additional experiments are needed to validate this speculation directly, for example by hepatocyte-specific CD44 depletion or inhibition in age-associated HCC models^41^.

We note several limitations of this study. First, our findings apply only to male sex. Aged female livers did not show increased hepatocyte CD44 expression (Fig. S1E), indicating sex-specific regulation of TICs, which warrants further study. These sex differences are consistent with epidemiologic and experimental evidence that male sex is a risk factor for HCC and suggest that sex hormones or sex-biased inflammatory signaling may modulate hepatocyte *Cd44* expression and functions in aged livers^42^. Second, the earliest timepoint at which *Cd44*-expressing hepatocytes appear during aging has not been defined. Third, bulk RNA sequencing of *Cd44* knockout liver does not isolate hepatocyte-intrinsic from non-parenchymal transcriptional changes. Fourth, the study does not establish direct causality between hepatocyte CD44 expression and T cell dysfunction. Finally, the study lacks human validation. All these points should be addressed in future studies.

The role of CD44 in a normal aging liver has been unclear, yet anti-CD44 strategies have been suggested to treat cancer^43^. This study provides evidence for a pathophysiological role of CD44 in aged livers where CD44 expressing hepatocytes activate the IL6/JAK/STAT3 pathway, generating a local immunosuppressive milieu with cytokine-deficient T cells. This may allow injured hepatocytes to persist and undergo transformation in the pre-cancerous niche. These findings can direct pre-clinical studies to decrease risk of cancer in aged liver, including consideration of rational combination strategies that target CD44, restore T cell function and/or modulate IL6/JAK/STAT3 signaling. In summary, our work identifies hepatocyte CD44 as a key age-associated node coupling TICs to IL6/JAK/STAT3 activation and local T cell dysfunction, thereby linking features of the aging liver microenvironment to increased HCC risk.

## 4. Methods

### Animals

Animal experiments were approved by the Institutional Animal Care and use Committee (IACUC) at Sanford Burnham Prebys (SBP) Medical Discovery Institute (Protocol numbers 22-033, 22-034, 25-031). Young and old C57Bl6J mice were obtained from the NIA aging colony housed at Charles River Laboratories or SBP in-house mouse colony. Mice were allowed to acclimate for 2-3 weeks at SBP Animal Facility before initiating experimental procedure. Mice were housed under specific pathogen-free conditions at 21-24 °C, 30-70 % humidity and a 12-hour light-dark cycle with ad libitum access to water and food (Teklad 2018). All injections were performed by trained veterinary staff at the SBP Animal Facility. Mice were monitored daily by the veterinary staff at SBP Animal Facility. No tumors were anticipated from the treatments in this study. Animals were humanely euthanized at the first occurrence of any of the following predefined criteria: (i) visible body weight loss, (ii) hunched posture, (iii) impaired mobility or moribund state; as advised by SBP animal facility veterinary staff. Mice were euthanized by CO2 asphyxiation. All experimental mice were euthanized between 8 am to 10 am unless recommended by veterinary staff to perform euthanasia as a humane endpoint. Any humane endpoint euthanasia was done immediately upon advice by the SBP veterinary staff. Young mice are defined as 4-8 months of age and old mice are defined as 18-25 months of age at the time of tissue collection. For adoptive transfer experiments, the AAALAC accreditation number is 000503. P14 C57Bl6J mice were obtained from Hans Peter Pircher^35, 36^. All P14 mice were housed at Salk Animal Facility as per ethical guidelines.

### AAV-mediated hepatocyte-specific CD44 knockout

Recombinant AAV8-TBG-saCAS9-U6-sgCd44g2 and AAV8-TBG-saCAS9-U6-sgCd44g4 were used to knockout Cd44 in old mice. For comparisons, recombinant AAV8-TBG-saCAS9-U6-sgRosa26 was used to knockout Rosa26 region as control. Virus suspension was prepared and injected at a dose of 3.33 x 10^11^ viral genomes per mouse in 100 μL sterile PBS retro-orbitally. The experiment was done in two independent cohorts. The end point of two months for this experiment was determined from results obtained from cohort 1. Liver tissues were either flash-frozen in liquid nitrogen or embedded in OCT (Tissue-Tek 4583) by rapid cooling or fixed in 10 % Neutral Buffered Formalin (epredia) for histology and immunofluorescence staining. Hepatocytes were isolated from the L lobe for flow cytometry analysis. Genomic DNA was extracted from about 50 mg frozen liver tissue using Monarch Genomic DNA Purification kit (New England Biolabs, Ref T3010S) and PCR was performed using 2X Taq PCR Master Mix (APExBIO, Ref K1034), 100 ng of gDNA and 1 μL of 100 μM primers to amplify the target region. The PCR products were purified QIAquick PCR Purification Kit (QIAGEN, Ref 28104) and submitted for Sanger sequencing. Using either the forward or the reverse primer. Indel frequency was quantified using ICE analysis as previously described^44^ (Synthego ICE v3.0). The CAS9 guide RNA sequences are:

i. sgCd44g2: caccgCAAATGAAGTTGGCCCTGAGC
ii. sgCd44g4: caccgTCACAGTGCGGGAACTCCCCC
iii. sgRosa26g1: caccGCTCGATGGAAAATACTCCGAG

The primer sequences used to amplify the guide RNA-targeted region in the genomic DNA are as follows.

i. sgRosa26g1. P_Rosa26_Fdw_1: CTAGAAGATGGGCGGGAGTCT, P_Rosa26_Rev_1: TCTAGGG GTTGGATAAGCCAGT.
ii. sgCd44g2. P_cd44_Fwd_2: ATGCAAAAGCCTCTAACATCTCC, P_cd44_Rev_2: CTACAGGAGCA CATTAGGCCA.
iii. sgCd44g4. P_cd44_Fwd_4: GGATTTCTACCCGCACGGTAT, P_cd44_Rev_4: CCTTTCACATGGA CGCTCTTT.

### Splenocyte isolation and T cell activation

Spleen was collected from 4–6-week-old P14 male mouse in RPMI 1640 (Gibco) in ice. Spleen was mashed on a 40 μm strainer using a 3 mL syringe plunger while passing ice-cold RPMI 1640 with 10 % FBS (Gibco). The flow through was collected on ice. The suspension was centrifuged at 420xg for 4 minutes at 4 °C followed by red blood cell lysis with ACK lysis buffer (KD Medical) for 3 minutes at room temperature. The lysis was neutralized with RPMI 1640 with 10 % FBS solution and centrifuged at 420xg for 4 minutes at 4 °C. The cells were counted with trypan blue (Sigma Aldrich) and plated at 1 million cells per well in a 12-well plate. T cells were stimulated with GP33 (GenScript Cat# RP20257, 1 μg/mL) and GP66 (1 μg/mL) in RPMI + 10 % FBS at 37 °C. The T cells were allowed to activate and expand for 48 hours at 37 °C and 5 % CO2. At the end of 48 hours, cells were counted with trypan blue and resuspended in RPMI with 10 % FBS at 1 million cells per 100 μL.

### Adoptive P14 T cell transfer

Recombinant AAV8-TBG-GP33DYDDK80 was used to express LCMV antigen GP33 in a hepatocyte specific manner. Recombinant AAV8-TBG-MCS (multiple cloning site, empty vector control) was used as control virus. Viral suspension was prepared at a dose of 1 x 10^6^ viral genomes per mouse in 100 μL and injected retro-orbitally. Experimental time point of 2 weeks for expression of GP33 was determined from previously literature^25^. After 2 weeks of AAV8 infection, ex vivo activated P14 T cells were injected at 1 million cells per 100 μL sterile T cell media. After 5 days of adoptive T cell transfer, mice were harvested. Livers were collected for P14 T cell analysis and IFNγ staining was done by flow cytometry.

### AAV8 generation

Viruses were obtained commercially from the Salk Viral Vector Core or the in-house SBP Functional Genomics Core. Briefly, HEK 293t cells were co-transfected with the AAV vector plasmid, RepCap8 expressing plasmid and the Helper plasmid (Cell Biolabs) using polyethylenimine (Polysciences). After 96 hours of transfection, both cells and the 96-hours media was collected for purifying the virus. Cells were lysed by sonication and treated with Benzonase (Millipore Sigma). The culture media was precipitated using polyethylene glycol. Supernatants were further subjected to iodixanol gradient ultracentrifugation to obtain an AAV fraction. The AAV fraction was then buffer exchanged with sterile PBS, aliquoted and stored at −80 °C. Viral titers were determined by SyBR Green qPCR using vector-specific primers.

### saCAS9 guide RNA design and validation

Guide RNAs were designed using CRISPick (Broad Institute) using the mouse reference genome GRVm38 (Ensembl v102). Guide RNAs targeting *Cd44* were designed, annealed and cloned into the vector pX602-AAV-TBG::NLS-SaCas9-NLS-HA-OLLAS-bGHpA;U6::BsaI-sgRNA (Addgene, 61593) as previously described^38^. Plasmids were transformed into Competent *E. coli* (NEB #C2988) and successful insertion was confirmed by Sanger sequencing (Genewiz) using universal U6 primer (5’-GACTATCATATGCTTACCGT-3’). Specificity of guides was first validated in vitro using AML12 and targeted deletion was confirmed by assessing cuts at the *Cd44* region of the genomic DNA. Genomic DNA was extracted using Monarch Genomic DNA Purification kit (New England Biolabs, Ref T3010S) and PCR was performed using primers flanking the target region (less than equal to 500 bp amplicon). The PCR was performed using 2X Taq PCR Master Mix (APExBIO, Ref K1034), 100 ng of gDNA and 1 μl of 100 μM primers. PCR products were then purified using QIAquick PCR Purification Kit (QIAGEN, Ref 28104) and submitted for Sanger sequencing using either the forward or the reverse primer. Indel frequency was quantified using ICE analysis (Synthego ICE v3.0 [04/15/2024]) as previously described^44^.

### Hepatocyte isolation for flow cytometry

For flow cytometry, hepatocytes were isolated from the L-lobe using Miltenyi Mouse Liver Perfusion Kit following manufacturer’s instructions (Miltenyi Biotec, #130-128-030). Mice were euthanized using CO2 asphyxiation and the L-lobe was extracted as early as possible and avoiding damage. The L-lobe was placed in chilled tissue storage solution (Miltenyi Biotec, #130-100-008) until all the mice were collected. After collection, L-lobe was placed on the Miltenyi perfuser (Miltenyi Biotec # 130-128-151) and perfused using Miltenyi Tissue Dissociator (Miltenyi Biotec #130-096-427) to obtain digested L-lobe and hepatocyte suspension. Hepatocytes were always kept on ice. The hepatocytes were counted with trypan blue, stained with a fixable viability dye (eBioscience #65-0865-14) and fixed (BioLegend FluoroFix Buffer #422101). The hepatocytes were blocked with anti-CD16/CD32 (1:250, TruStain FcX™ PLUS anti-mouse CD16/32, BioLegend 1#56603) antibody and stained with CD44 antibody (Clone IM7, BioLegend #103009; BioLegend #103007) and CD45 (Clone 30-F11, BioLegend #103147). Events were acquired within 24 hours using Cytek Aurora 5L spectral flow cytometer. Data acquired was unmixed using SpectroFlo (Cytek Biosciences). Data was further analyzed using FlowJo v10 or FlowJo v11 to obtain representative flow plots and quantitation.

### Liver immune cell isolation and flow cytometry

Whole liver was dissected out and placed on a 40 or 70 μm cell strainer mounted on 50 mL tube. Single cell suspension was obtained by mashing using 3 mL syringe plunger while passing 10 % FBS in RPMI. Liver samples were centrifuged at 60 r.c.f., 4 °C for 2 minutes with no brake to pellet hepatocytes. The supernatant was collected and centrifuged at 420 r.c.f, 4 °C for 4 minutes. Immune cells were enriched with 40 % Percoll (Cytiva) in HBSS. The pellet was resuspended in ACK lysis buffer (KD Medical) for red blood cell lysis. The lysis buffer was neutralized and removed by centrifugation. Cells were then counted with trypan blue and hemocytometer. A FoxP3 transcription factor staining kit (eBioscience Cat# 00-5523-00) was used for intracellular staining. Viability staining was performed using LIVE/DEAD fixable red stain (1:1000 in FACS buffer, Invitrogen) for 15 minutes at room temperature. Suspensions were then pelleted and resuspended in anti-CD16/32 antibodies (1:500, TruStain FcX™ PLUS anti-mouse CD16/32, BioLegend #56603) to block non-specific binding to Fc receptors. Cells were incubated with the indicated with surface antibodies for Thy1.1 (Clone OX-7, BD #612837), CD8a (clone 53-6.7, BD #563786) and PD-1 (Clone 29F.1A12, BioLegend #135225) antibodies for 30 minutes at 4 °C. Antibodies against IFN-γ (Clone XMG1.2; BioLegend #505829; BioLegend #505809) were diluted in 1X permeabilization buffer and added for 45 minutes at 4 °C. For cytokine staining, cells were stimulated with PMA (final concentration of 1 μg/mL) and ionomycin (Cell Signaling; final concentration of 1 μg/mL) for 4 h at 37 °C in the presence of brefeldin A (GolgiPlug, BD Biosciences; final concentration of 1 µg/mL) to block cytokine export from the golgi apparatus. 2% paraformaldehyde (PFA) was used to fix the cells after staining. Cells were resuspended in 100 μL 1X PBS and events were collected on the FACSymphony A3 5-laser flow cytometer (BD Biosciences) within 48 hours of fixing. Data were analyzed using FlowJo (v.10, BD Biosciences).

### Fluorescence-activated cell sorter

For 10X scRNAseq, immune cells were isolated from the livers as described above. Immune cells from 7 young and 7 old mice were combined to enrich for adoptively transferred P14 CD8+ T cells. First, T cells were enriched using Invitrogen T cell enrichment kit (Invitrogen 8804-6822) in low-bind FACS tubes (Falcon). The enriched T cells were then stained for the congenic marker Thy1.1 (Clone HIS51, eBioScience # 12-0900-81), PD-1 (Clone 29F.1A12, BioLegend #135209), CD8 (Clone 53-6.7, BioLegend # 100726), DAPI (0.5-1 μg/μL). Antigen-specific response was confirmed by anti-PD1 response observed during sorting. Thy1.1+ CD8a+ cells were sorted out using flow cytometer at the SBP flow cytometry core on a SORP FACSAria IIu (BD Biosciences) with 100 μm nozzle, collected in 0.05% BSA in PBS (Miltenyi). Cells were counted for viability using AO/PI dye in K2 Cellometer (Revity) and then processed for single cell RNA sequencing.

### Plasmids

The following plasmids were used in this study:

i. pX602-AAV-TBG-SaCas9-U6-Rosa26_g1
ii. pX602-AAV-TBG-SaCas9-U6-sgCd44 (gRNA2)
iii. pX602-AAV-TBG-SaCas9-U6-sgCd44 (gRNA4)
iv. pAAV[Exp]-TBG>{GP33_DYDDK_80}:WPRE (produced by Vector Builder)
v. AAV-TBG-MCS_108bp (produced by Vector Builder)

### Liver single nuclei multiome

Raw sequencing reads were processed using cellranger arc (v2.0.2). Ambient RNAs were removed using CellBender (v0.3.0)^45^. Cells marked as doublets by DoubletFinder (2.0.4)^46^ were removed. Additional data processing was performed using Seurat (v5.2.1)^47^ and Signac (v1.12.0)^48^. We also discarded cells with >10% mitochondrial reads, <200 genes detected, <1000 ATAC reads, or ATAC enrichment at TSS < 1. The samples were then merged, log-normalized, and integrated using Harmony. After cell clustering by Seurat, initial cell type annotation for each cluster was performed based on canonical marker genes. For each initial cell type, we performed sub clustering and checked the expression of marker genes for each subclusters. Subclusters expressing multiple marker genes were removed. Hepatocytes were considered *Cd44*+ if their normalized expression of *Cd44* was greater than 0. Differentially expressed genes between *Cd44*+ and *Cd44*− hepatocytes were performed using the function FindMarkers in the Seurat package with the default parameters except the following parameters: test.use = “Mast”, min.pct = 0.1, logfc.threshold = 0.1. For the ATAC analysis, gene activity and transcription factor activity by chromVar^49^ were calculated using Signac^48^. GO analysis was done using the function enrichGO() in the package clusterProfiler (v4.6.2). Significantly enriched GO terms were defined by adjusted p-value<0.05.

### RNA sequencing and analysis

RNA was analyzed using Bioanalyzer. Libraries were prepared and sequenced by the SBP Genomics core. PolyA RNA isolation and library preparation was performed with the Watchmaker mRNA Library Prep Kit (Watchmaker Genomics: 7BK0001), xGEN Stubby adaptors (IDT, Cat: 336338420) and xGEN 10nt UDIs (IDT, Cat: 336338436). Libraries were sequenced with the Element Biosciences AVITI Sequencing platform using the AVITI 2×75 High Output Cloudbreak Freestyle Kit (Element Biosciences, Cat: 860-00015). For all experiments, raw reads were quality-checked with FastQC v0.11.8. Experiments containing adaptor sequences were trimmed with Trim Galore v0.4.4_dev. Reads were aligned to the mm10 reference genome with STAR aligner v2.5.3a^50^ and converted to gene counts with HOMER’s analyzeRepeats.pl script^51^. Gene counts were normalized and queried for differential expression using DESeq2 v1.30.0^52^. For each pairwise comparison, genes with fewer than 10 total raw counts across all samples were discarded prior to normalization, and genes with an absolute log2 fold change > 0.5 and an FDR-corrected p-value ≤ 0.05 were pulled as significant. Pathways were queried for treatment-specific functional enrichment using gene set enrichment analysis (GSEA) in WebGestaltR v0.4.4^53^ using rank-ordered log2foldchange values.

### CosMx analysis

The M lobe (Median lobe) of each mouse liver (n = 12) was excised immediately after euthanasia; one mouse (ID 1177) was first cardiac-perfused with ice-cold PBS and zinc-based fixative (Z-fix, Cancer Diagnostics, Inc.). All excised lobes were incubated in fresh Z-fix for 48 hours at room temperature, then transferred to 70 % ethanol and stored at 4 °C. Livers were processed and embedded in paraffin according to CosMx recommended guidelines. The blocks were sectioned on a microtome to obtain 5 µm scrolls, which were floated on an RNase-free, molecular-grade water bath and collected onto slides. Slides were then processed for CosMx spatial transcriptomic by Bruker Spatial Biology, Seattle WA. Flat files for each CosMx slide were processed using Seurat (v4.9.9.9050). The individual tissue samples from each slide were split and to be treated as individual samples. The FOVs for each tissue (nine/tissue) were combined to form one large FOV/tissue. Cells in the top and bottom 15th percentile for number of features and with counts less than 20 were removed. The samples were then re-merged, normalized using SCTransform (v0.3.5) with a clip range of −10 to 10, and integrated using Harmony (v1.2.3). Each cluster, as determined by Seurat, was further subclustered (resolution 0.1 to 0.3). Cell type annotation was first checked using CellKB^54^ after filtering for only “liver” studies. The top five markers, sorted by log2 fold change were used as input genes for CellKB. Cell type annotation was further confirmed using known markers for liver cell types (Immune Cells: *Ptprc*; B Cells: *Cd79a*; Cholangiocytes: *Spp1*, *Krt19*, *Epcam*, *Sox9*; Hepatocytes: *Apoa1*, *Glul*, *Serpina1a*, *Apoe*; HSC: *Dcn*, *Bmp5*, *Col1a1*, *Col1a2*; Kupffer Cells: *Cd5l*, *Cd74*, *C1qa*, *C1qb*; LSEC: *Lyve1*; NK Cells: *Stat4*, *Nkg7*; T Cells: *Cd2*, *Cd3d*, *Cd3g*). Hepatocytes were considered *Cd44*+ if their normalized expression of *Cd44* was greater than 0. Neighborhoods were defined as the 8 nearest neighbors (KNN) of *Cd44*+ hepatocytes.

### MERSCOPE analysis

MERSCOPE images were segmented using the Vizgen Post-processing Tool (VPT, v1.3.0) with Cellpose (models cyto2 and nuclei, 2D). DAPI and PolyT channels were normalized by CLAHE before segmentation. Cell and nuclear masks were generated on the central z-plane (z = 3) and harmonized into final cell boundaries using default VPT fusion parameters. Cells containing greater than 100 transcripts and 10 unique genes were retained after quality filtering. Samples were SC-Transformed, integrated with harmony, and UMAP-clustered in Seurat v5.0.1.9001. Individual Seurat clusters were annotated using a panel of liver cell typing genes, in addition to supplying the top 20 most cluster-specific genes to CellKB. Neighborhoods were defined as the 8 nearest neighbors (KNN) of *Cd44*+ hepatocytes.

### P14 immune single cell analysis

For each sample, shallow and deep sequencing fastq files were processed using Cell Ranger v8.0.1^55^. Filtered matrix files were processed using Seurat v4.3.0.1^56^. Cells with nFeature less than 200, nFeature greater than 4,000, or percent mitochondrial DNA greater than 5 % were removed from each sample. Samples were normalized using SCTransform^57^ and integrated following Seurat’s default pipeline using Harmony v1.2.0^58^. Differentially expressed genes were calculated using FindMarkers and the Wilcoxon test (p < 0.05, abs (log2FC) > 0). Gene set enrichment analysis (GSEA) was performed using the WebGestaltR package (version 0.4.6) in R with the MSigDB mouse collections. Ranked gene lists were generated by ordering all genes according to their signed fold change values obtained from the differential expression analysis of all old cells vs. all young cells. Normalized enrichment scores (NES) and FDR-adjusted p values were used to determine significantly enriched pathways.

### Ingenuity Pathway Analysis

Upstream regulator analysis on differentially expressed genes was performed using Qiagen’s Ingenuity Pathway Analysis software. The adjusted p value threshold for analysis was set at less than 0.05. Predicted upstream regulators were visualized as volcano plots (activation z-score vs –log10 p-value of overlap) using Matplotlib in JupyterLab 4.3.4.

### Statistics

The following statistical tests were done in the study. For Fig. 1A, adjusted p values were determined from DEG analysis with DESeq2. For Figs. 1B and 1C, p value was determined using Wilcoxon rank sum test. For Figs. 1D and S1B, p values were determined by t test between log-normalized *Cd44* expression in young and old hepatocutes for all cells expressing *Cd44* (*Cd44*-cells were excluded). Figures show data from same mice split by H2O2 treated and untreated slides. For Figs. 1E and 3A, p values were determined by unpaired t test with Welch’s correction in Prism 10.6.1. For Figs. 2A and S2A, p values were calculated by MAST method in Seurat package. For Fig. 2F, p value was determined using Wilcoxon rank sum test after calculating motif activity for all cells using ChromVar comparing *Cd44*+ hepatocytes with *Cd44*− hepatocytes. For Fig. 3B, p value was determined by Wilcoxon test (p < 0.05, abs(log2FC) > 0). For Figs. 4B, 4C and S2D; GSEA was performed on the rank-ordered fold changes from each comparison. For Fig. S3B data were analyzed using Kruskal–Wallis test and multiple comparisons were corrected using the two-stage linear step-up procedure of Benjamini, Krieger and Yekutieli in Prism 10.6.1. For Fig. S3C p value was determined using unpaired t test in Prism 10.6.1. For Fig. S3E differential expression analysis between all P14 CD8+ cells recovered from old versus young was done using Seurat’s FindMarkers function with Wilcoxon text with Benjamni-Hochberg correction to report adjusted p values. For Fig. S4E, p values were determined using Ordinary one-way ANOVA with uncorrected Fischer’s LSD in Prism 10.6.1. P value of less than 0.05 was considered significant.

## 5. Data Availability

All RNA-seq and spatial transcriptomic datasets generated in this study have been deposited in the Gene Expression Omnibus (GEO) under the following accession numbers. GSE313646: RNA sequencing of hepatocytes isolated from young and old mouse livers. GSE310240: Bulk RNA-seq of liver tissue from hepatocyte-specific Cd44 knockout and Rosa26 control mice. GSE304160: CosMx spatial transcriptomics of young and old mouse liver. Single nuclei multiome from young and old mouse livers is available at (https://data.sennetconsortium.org/search?size=n_20_n&sort%5B0%5D%5Bfield%5D=last_modified_timestamp&sort%5B0%5D%5Bdirection%5D=desc). GSE309965: Single cell RNA sequence data of P14 CD8+ T cells recovered from young and old livers.

## Acknowledgements

Work in the lab of PDA was supported by P01 AG073084, P01 AG031862 and U54 AG079758. Authors would like to acknowledge the Animal Facility Core (Adriana Charbono and Buddy Charbono); Histology Core (Gia Guillermina and Monical Sevilla); Flow Cytometry Core at Sanford Burnham Prebys Medical Discovery Institute and Sanford Burnham Prebys Functional Genomics Core. Flow cytometry is supported by (Benji Portillo/Yoav Altman and) the SBP Flow Cytometry Shared Resource (RRID:SCR_014854) with funding from NCI CCSG P30CA030199. This study includes data generated with the Cytek Aurora flow cytometer funded by Shared Instrumentation Grant S10OD032325. Functional Genomics Core is supported by Sanford Burnahm Prebys NCI Cancer Center Support Grant P30 CA030199 and Shared Instrumentation Grant S10 OD036254.

## Author Contributions

PDA and AG conceptualized the study. PDA secured funding for the study. AG designed and performed the experiments. AG wrote the manuscript and prepared the figures. YL, KL, GE, CM, QY, ES, SM, AW, BR, KY and AW performed bioinformatic analysis and plot generation. FAH, ML, ST, SK and PDA provided reagents. SK and PDA conceptualized T cell experiments. RA, MGT, SY, AH, AR, JP, BP and AD provided technical help.

---

**Figure S1.**
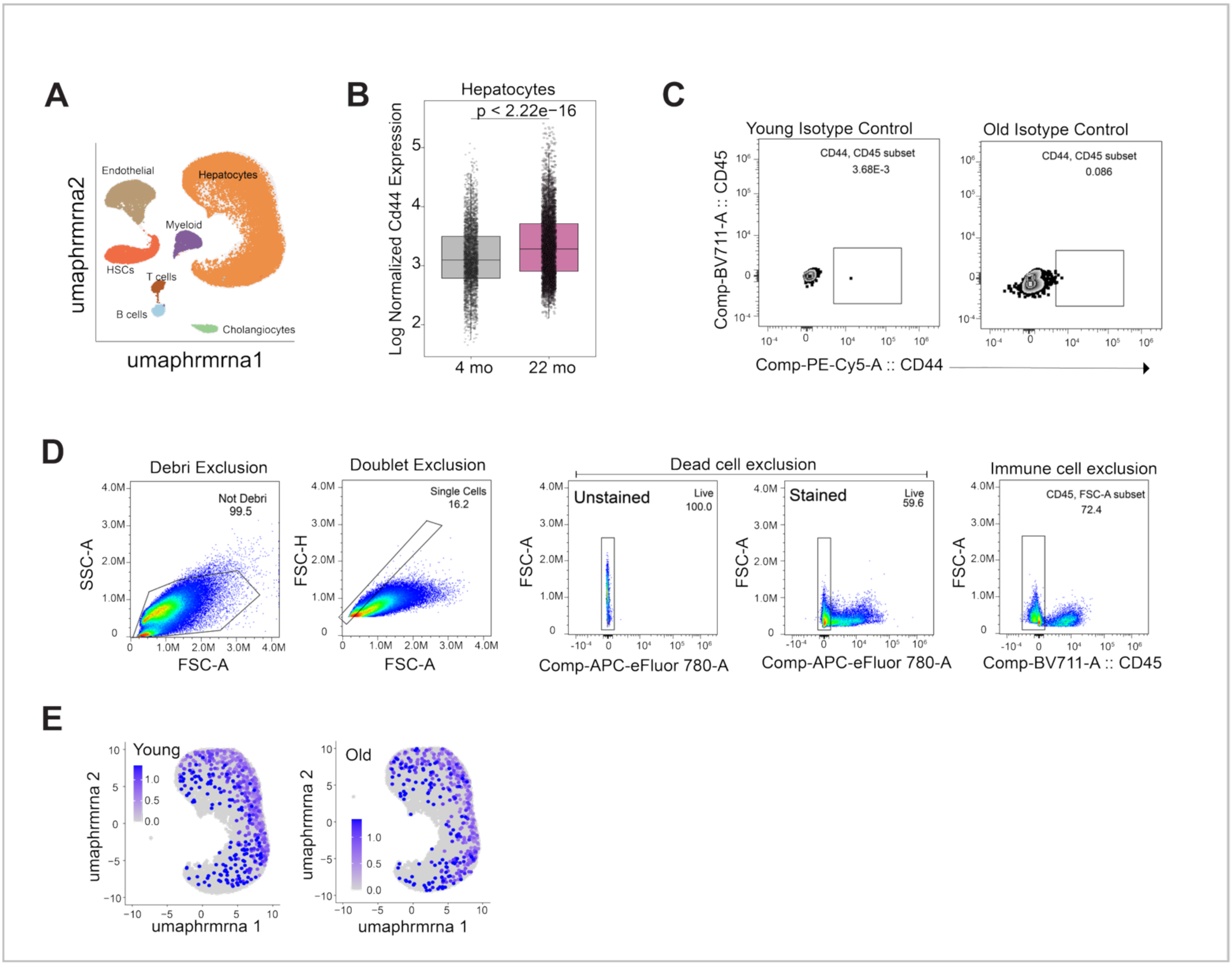
Abundance of CD44-expressing hepatocytes increases in aged mouse liver. **A.** Uniform Manifold Approximation and Projection (UMAP) of liver nuclei (10x multiome) from young and old C57BL/6J male mice, annotated by cell type (Endothelial, Hepatocytes, Myeloid, Hepatic stellate cells, T cells, B cells, Cholangiocytes). **B.** Box plot shows distribution of *Cd44* expression per hepatocyte with age. Y-axis represents log-normalized expression of *Cd44* in hepatocytes. **C.** Flow cytometry analysis of hepatocytes from young and old mouse livers. L-lobes were perfused, and hepatocytes were purified using Miltenyi perfusion kit. Flow plots show staining with isotype control antibodies in young and old isolated hepatocytes. **D.** Gating strategy for Fig. 1E and S1C. **E.** Single nucleus multiome profiling of young and old female mouse livers. UMAP shows *Cd44* transcript expression in hepatocyte nuclei from young (n = 3) and old (n = 3) female livers, each dot represents a nucleus.

**Figure S2.**
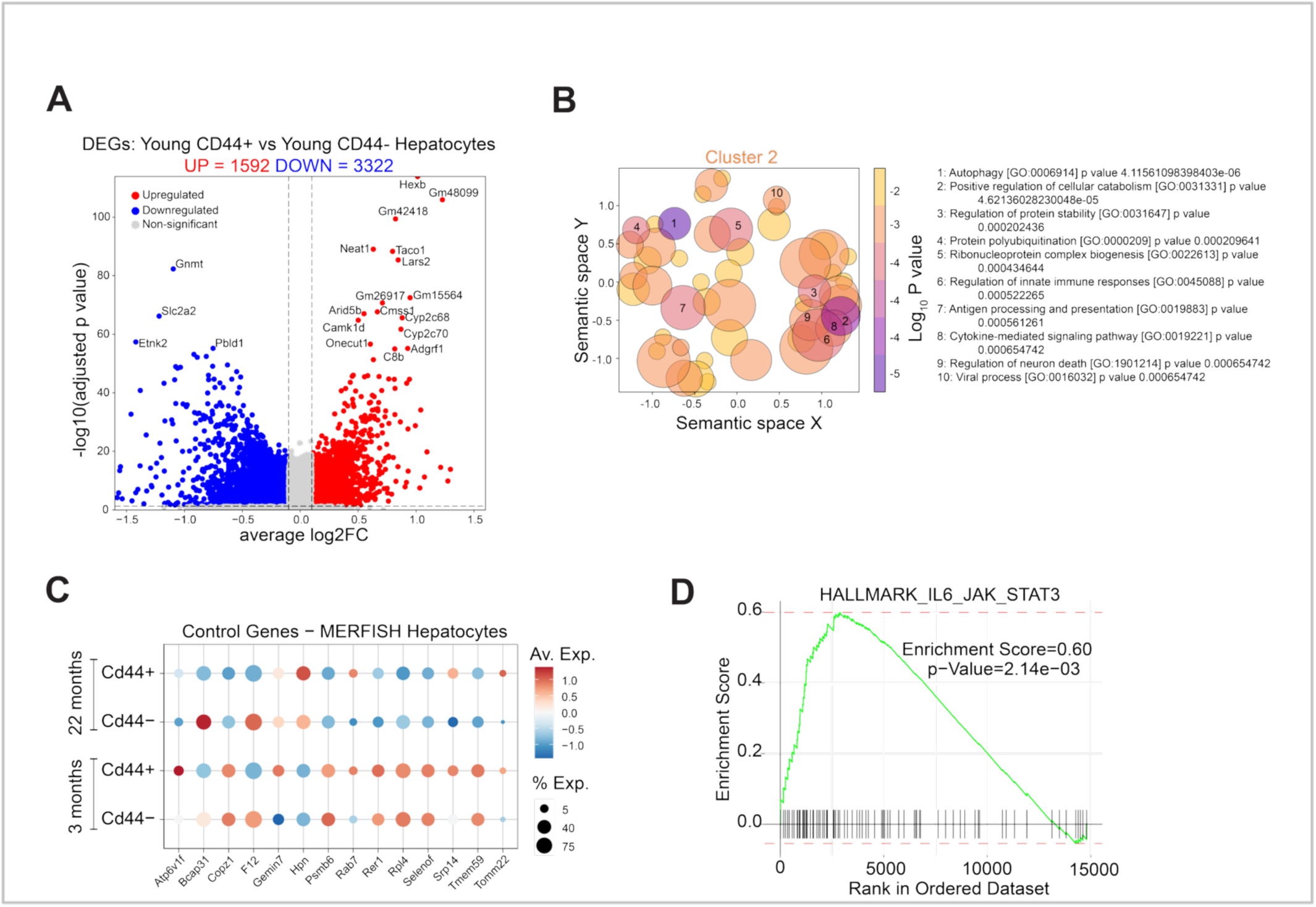
*Cd44* expressing hepatocytes in aged liver exhibit an immune modulatory transcriptome. **A.** Volcano plot depicting DEGs between *Cd44*+ versus *Cd44*-hepatocytes in young male livers. 1592 genes are upregulated and 3322 genes are downregulated in young *Cd44*+ hepatocytes compared to young *Cd44-* hepatocytes. (Log2FoldChange cutoff of 0.1 and −0.1 respectively, and adjusted p-value ≤ 0.05). **B.** GO term enrichment analysis of genes from Cluster 2 in Fig. 2D. GO terms are displayed in semantic space by GoFigure to reduce redundancy. Top 10 significant enriched terms labelled. **C.** Expression of control genes by MERSCOPE in hepatocytes defined as *Cd44*+ and *Cd44*-in young and old livers. Color indicates average expression and size of dot indicates percent of cells expressing cells. **D.** Gene Set Enrichment Analysis (GSEA) for IL-6/JAK/STAT3 signaling pathway in hepatocytes isolated from old livers compared to young livers^25^. Enrichment score = 0.6 in old hepatocytes compared to young hepatocytes, FDR = 2.14e−03.

**Figure S3.**
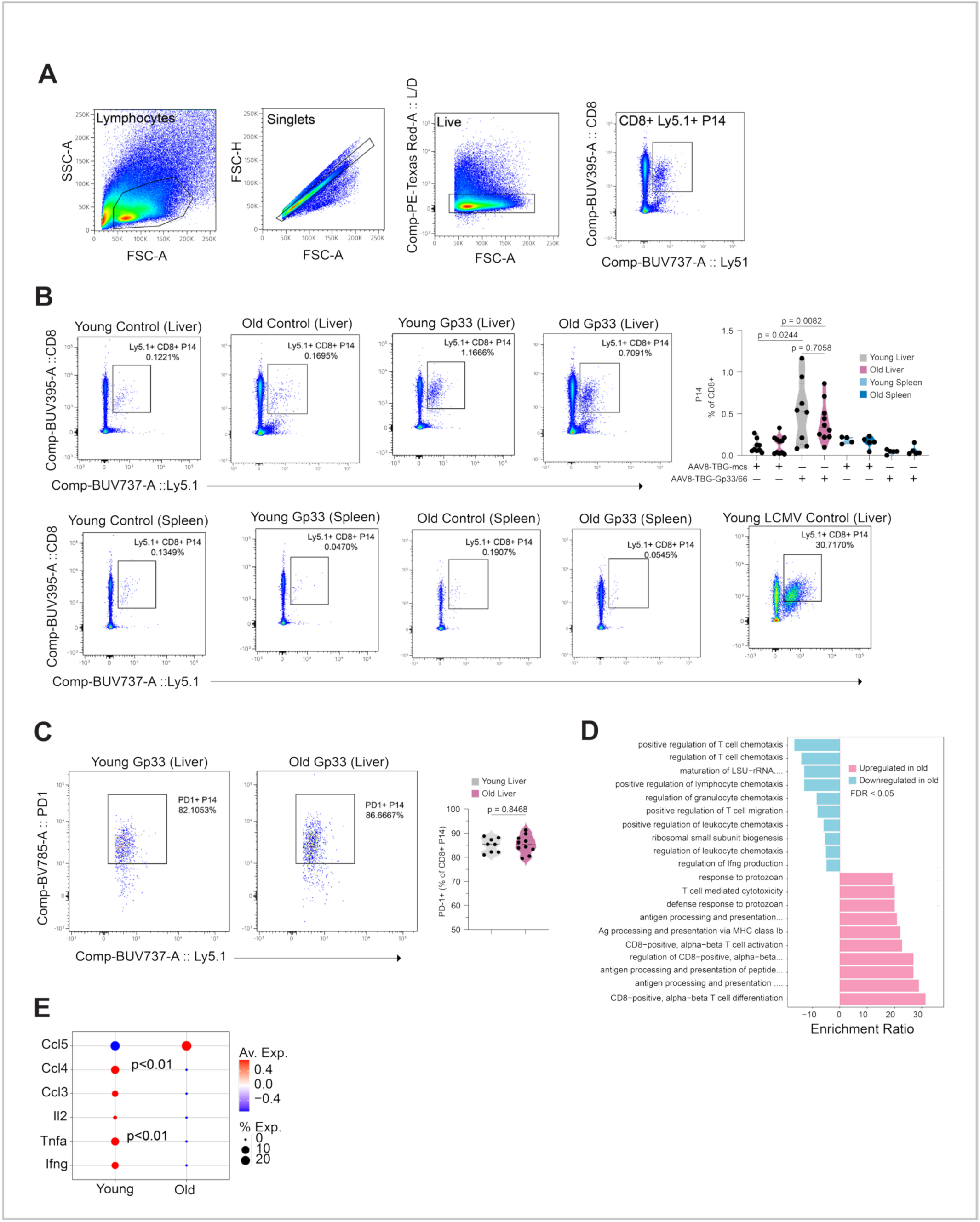
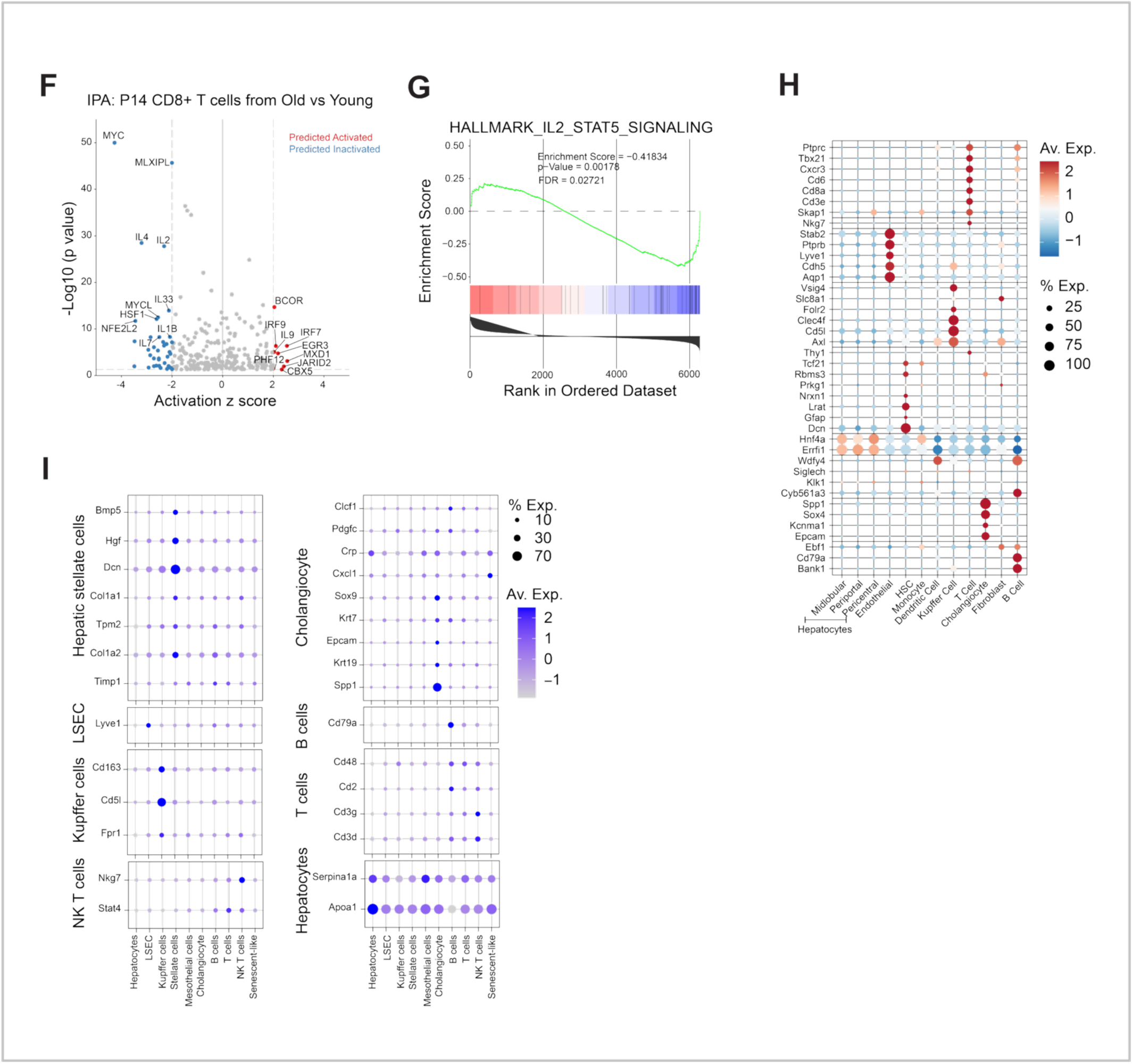
Aged livers harbor T cells with markers of inactivation. **A.** Gating strategy for analyzing Ly5.1+ P14 CD8+ T cells by flow cytometry. CD8+ Ly5.1+ P14 cells were gated on live (negative for PE-Texas Red) and single cells. Lymphocytes were first gated by forward and side scatter, followed by singlets (FSC-A vs. FSC-H) and live cells (PE–Texas Red–negative). From this live, single lymphocyte population, CD8⁺ Ly5.1⁺ P14 T cells were gated for analysis. **B.** Representative flow plots depicting staining for CD8 versus Ly5.1 in cell suspension of young and old livers and spleens of mice infected with AAV-TBG-mcs (control) and AAV-TBG-GP33 (antigen), and liver of young LCMV infected positive control. Violin plot quantifies the percentage of CD8+ Ly5.1+ cells in the total CD8+ T cell population recovered from young and aged, control or antigen infected livers; each dot represents one mouse (n = 9 young liver (control), n = 11 old liver (control), n = 8 young liver (antigen), n = 10 old liver (antigen), n = 4 young spleen (control), n = 5 old spleen (control), n = 5 young and old spleen (antigen). p values indicated for relevant comparisons in the plot. Data were analyzed using Kruskal–Wallis test and multiple comparisons were corrected using the two-stage linear step-up procedure of Benjamini, Krieger and Yekutieli. **C.** Representative flow plots show Ly5.1 versus PD-1 staining gated on CD8+ P14 population. Violin plot quantifies the PD-1+ as a percent of the CD8+ P14 population recovered from young and aged livers; each dot represents one mouse (n = 8 young, n = 10 old); p value was calculated by Unpaired t test (p = 0.8468, non-significant). **D.** GO term enrichment analysis of upregulated and downregulated DEGs (Significant ( p < 0.05) with a log2FC of more than 1 or less than −1) from single cell RNA-seq of P14 CD8+ T cells from old recipient livers compared to young recipient livers. Cells pooled from n = 7 young and n = 7 old mouse livers. **E.** Dot plot showing expression of effector cytokines, *Il2* and chemokines from single cell RNA-seq of P14 CD8+ T cells recovered from young and old recipient livers. Color denotes average expression level and size of dot indicates percent of cells expressing the transcript. Significant p values indicated in the plot. Adjusted p values were calculated by Benjamni-Hochberg test. **F.** Ingenuity pathway analysis of DEGs comparing P14 CD8+ T cells recovered from young and old recipient livers shown in Fig. 3B. Volcano plot depicts top upstream predicted activated (Activation z-score ≥ 2) and predicted inactivated (Activation z-score ≤ −2) regulators in P14 CD8+ T cells recovered from old livers. **G.** GSEA for IL2/STAT5 signaling pathway in P14 CD8+ T cells recovered from old recipient livers compared to young livers. Enrichment score = −0.41834 in P14 cells recovered from old compared to young, FDR = 0.02721. **I.** and **J.** Cell type annotation marker expression plot for MERSCOPE and CosMx respectively.

**Figure S4.**
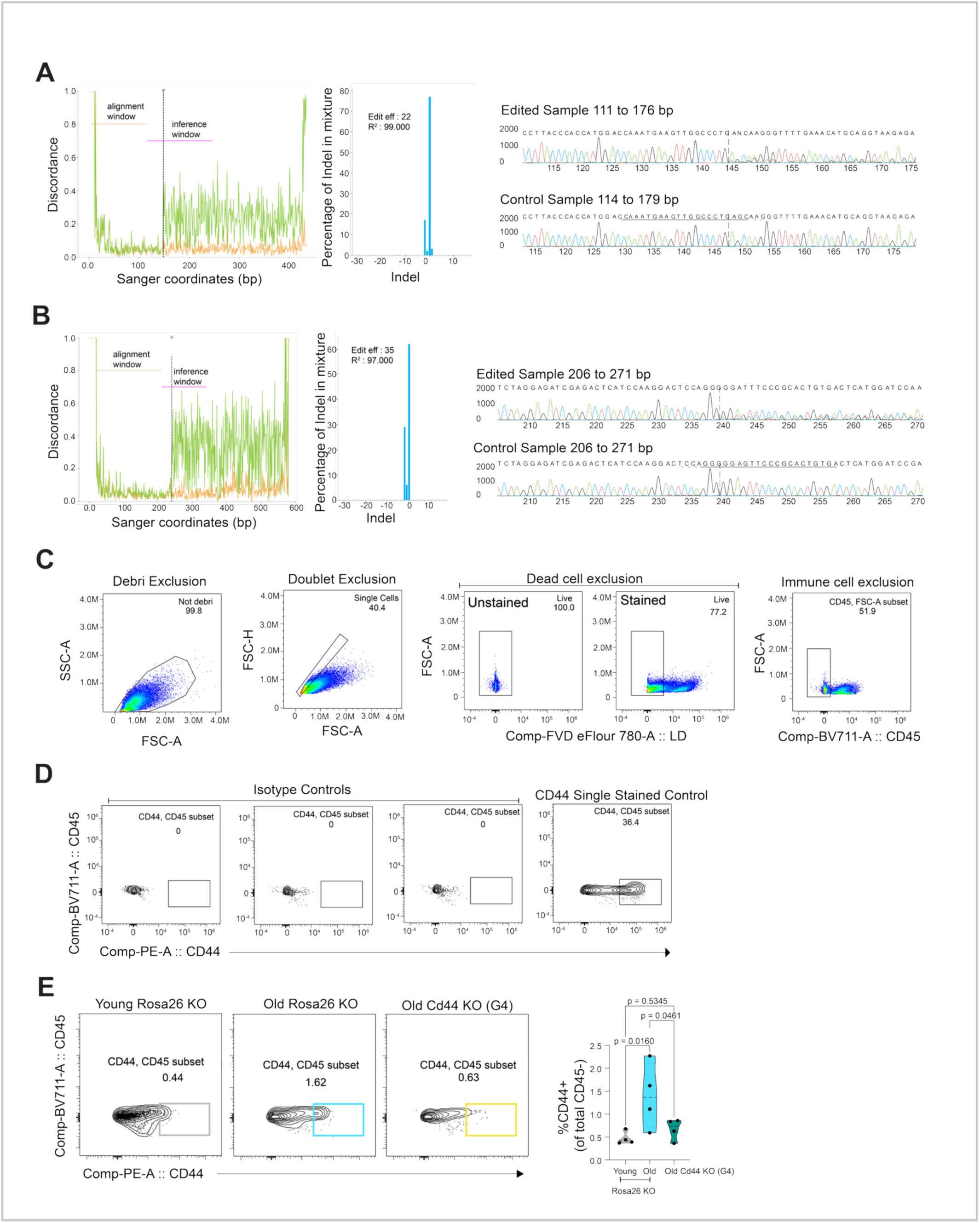
Knock out of hepatocyte *Cd44* attenuates immune modulatory gene signatures in aged mouse livers. **A.** and **B.** Left: *Cd44* knockout in old livers using sgRNAg2 (A) and sgRNAg4 (B). Plot depicts discordance in the *Cd44* gene at the inference window. Middle: Insertions/deletions in *Cd44* gene are quantitated as Indel. Right: Corresponding edited sample sequence display with the control sample sequence. **C.** Flow cytometry analysis of CD44 in hepatocytes from young and old mouse livers. L-lobes were perfused, and hepatocytes were purified using Miltenyi perfusion kit. Flow plots show gating strategy for analyzing CD44 staining on hepatocytes. Hepatocytes were identified by first excluding debris and doublets, followed by gating on live cells and subsequently excluding CD45+ cells. **D.** Flow plots show staining with isotype controls of CD45-population from C for young and old isolated hepatocytes. CD44 single stained control was used for gating on CD44+ population. **E.** Representative flow plots show CD44 versus CD45 staining of CD45-cells from young Rosa26 control, old Rosa26 control and old Cd44 knockout with sgRNAg4. Corresponding violin plot quantifies the percentage of CD44+ population within the CD45 negative population by age; each dot represents one mouse (n = 4 young (Rosa26 control), n = 4 old (Rosa26 control), n = 4 old (Cd44 knockout). p values for all comparisons were determined by Ordinary one-way ANOVA with uncorrected Fischer’s LSD in Prism 10.6.1.

**Supplementary Data Table 1 (T1).** Differentially expressed genes (DEGs) in young *Cd44*+ hepatocytes: full DEG list comparing young *Cd44*+ versus young *Cd44*− hepatocytes (depicted in Fig. S2A). DEGs were identified using the MAST method applied to single-nucleus RNA-seq data.

**Supplementary Data Table 2 (T2).** Differentially expressed genes (DEGs) in old *Cd44*+ hepatocytes: full DEG list comparing old *Cd44*+ versus old *Cd44*− hepatocytes (Fig. 2A). DEGs were identified using the MAST method applied to single-nucleus RNA-seq data.

**Supplementary Data Table 3 (T3).** Gene Ontology (GO) enrichment for genes uniquely upregulated in old Cd44+ hepatocytes: complete list of GO terms derived from enrichment analysis of the 186 genes uniquely upregulated in old Cd44+ hepatocytes.

**Supplementary Data Table 4 (T4).** Gene Ontology (GO) enrichment for Cluster 2 genes: complete list of GO terms derived from enrichment analysis of genes in Cluster 2 (depicted in Fig. 2D).

**Supplementary Data Table 5 (T5).** Upstream regulator analysis for Cd44+ hepatocyte DEGs in old liver: complete list of predicted upstream regulators identified from DEGs comparing *Cd44*+ versus *Cd44*− hepatocytes in old liver.

**Supplementary Data Table 6 (T6).** Single cell RNA seq of P14 CD8+ T cells recovered from old livers: full list of DEGs for P14 CD8+ T cells recovered from GP33-expressing old livers in comparison to young livers. Cells were pooled from n = 7 young and n = 7 old mouse livers.

**Supplementary Data Table 7 (T7).** Upstream regulator analysis for P14 CD8+ T cells recovered from old livers: complete list of predicted upstream regulators identified from DEGs in P14 CD8+ T cells recovered from old livers compared to young livers.

